# Putaminal dopamine modulates movement motivation in Parkinson’s disease

**DOI:** 10.1101/2024.03.15.585221

**Authors:** Magdalena Banwinkler, Verena Dzialas, Lionel Rigoux, Adrian L. Asendorf, Hendrik Theis, Kathrin Giehl, Marc Tittgemeyer, Merle C. Hoenig, Thilo van Eimeren

**Author notes:** Correspondence to: Thilo van Eimeren, MD, FEAN, University Hospital Cologne, Department of Nuclear Medicine, Kerpener Str. 62, 50937 Cologne, Germany.

## Abstract

The relative inability to produce effortful movements (akinesia) is the most specific motor sign of Parkinson’s disease. The motor motivation hypothesis suggests that akinesia may not reflect a deficiency in motor control *per se*, but a deficiency in cost-benefit considerations for motor effort. For the first time, we investigated the quantitative effect of dopamine depletion on the motivation of motor effort in Parkinson’s disease.

A total of 21 patients with Parkinson’s disease and 26 healthy controls were included. An incentivized force task was used to capture the amount of effort participants were willing to invest for different monetary incentive levels and dopamine transporter depletion in the bilateral putamen was assessed.

Our results demonstrate that patients with Parkinson’s disease applied significantly less grip force than healthy controls, especially for low incentive levels. Congruously, decrease of motor effort with greater loss of putaminal dopaminergic terminals was most pronounced for low incentive levels. This signifies that putaminal dopamine is most critical to motor effort when the trade-off with the benefit is poor.

Taken together, we provide direct evidence that the reduction of effortful movements in Parkinson’s disease depends on motivation and that this effect is associated with putaminal dopaminergic degeneration.

## Introduction

Dopamine is known for its crucial role in motivated behavior, such as investing more physical effort or waiting for longer periods in return for larger rewards.^1–3^ Current frameworks suggest that this effect arises from dopamine’s ability to promote goal-directed behavior by attributing incentive salience to stimuli and overcoming effort costs.^4–6^

Parkinson’s disease (PD) is characterized by severe depletion of dopaminergic terminals in the putamen, and by the relative inability to produce effortful movements (i.e., akinesia).^2^ The recently proposed “*motor motivation hypothesis”* suggests that akinesia in PD may not reflect a deficit in direct motor control *per se*, but a deficiency in the cost-benefit consideration for motor effort.^7^ In other words, besides the role of dopamine in explicit reward-seeking behavior, dopamine may also provide the substrate for *“movement motivation”*.

An increasing body of literature substantiates this motor motivation hypothesis of PD. However, direct evidence of the link between dopaminergic degeneration and movement motivation is still missing. For example, studies have demonstrated that the willingness to invest physical effort is dependent on the medication status of patients with PD, such as that medicated patients invest more effort at higher speed compared to unmedicated patients.^8,9^ These results can potentially be explained by low dopamine levels in the unmedicated state, which lead to a shift in the cost-benefit evaluation of effort, and thus a deficiency in performing effortful movements to obtain a reward.^9–12^ Additionally, it has been reported that independent of the medication status, patients with PD invest significantly less effort than healthy controls, especially for low reward options.^13^ Since previous studies on incentive salience in PD are limited to general group (PD vs. HC) and medication (ON vs. OFF dopaminergic medication) effects, quantitative measures of baseline endogenous dopamine and associated changes in movement are needed.

Thus, we investigated the relationship between putaminal dopaminergic terminal loss (using DaT (dopamine transporter) SPECT, an established standard for assessing dopaminergic degeneration^14^) and performance in an incentivized grip-force task within a group of unmedicated patients with PD. We hypothesized that unmedicated patients with PD would invest significantly less effort compared to healthy controls (HC). Additionally, in the PD group, we anticipated that greater putaminal dopaminergic terminal loss would be parametrically associated with lower effort and that this deficit would be more evident for low incentives.

## Materials and methods

### Participants

21 patients with PD and 26 HC between the age of 49 and 77 years were included in the study. Participants were recruited at the University Hospital Cologne, through posters and flyers, and from a local register of healthy volunteers. Prior to inclusion, all participants were screened for eligibility. Exclusion criteria were left-handedness, cognitive deficits (Montreal Cognitive Assessment [MoCA] score of < 24), and depression (Geriatric Depression Scale [GDS] score of > 5) as well as any significant comorbidity. For patients with PD, additional inclusion criteria were early-stage PD (clinical symptoms < 3 years), diagnosis according to the Movement Disorder Society Clinical Diagnostic Criteria for PD,^15^ and an available DaT SPECT, acquired shortly before the behavioral assessment at the University Hospital Cologne.

### Study procedure

The study was approved by the Ethics Committee of the University Hospital Cologne and performed in accordance with the standards of the Declaration of Helsinki. All participants gave written informed consent and received monetary compensation for their participation. The study included collecting demographic information, evaluating motor symptoms using the Unified Parkinson’s Disease Rating Scale (UPDRS-III), and a computerized incentivized force task. DaT SPECTs of patients with PD, which were acquired at the Department of Nuclear Medicine (University Hospital Cologne) as part of the clinical routine, were used for the analysis. Most patients were unmedicated. Medicated patients were asked to pause their medication prior to their study visit. Long-acting dopamine agonists were discontinued for at least 72 and L-DOPA for 12 hours.

### Incentivized force task

We used a well-established paradigm, which enables to quantify the effort (i.e. grip force) participants are willing to invest for different monetary incentive levels. More details on the task can be found elsewhere.^8,16,17^

Briefly, a hand dynamometer (Vernier Go Direct, USB connected) was placed in the participant’s right hand and the maximum voluntarily grip force was assessed to adapt effort levels in the task to each participants’ individual strength. In the task, participants were instructed to maximize their total monetary payoff in two conditions: (A) win and (B) loss avoidance (Fig. 1). Each condition comprised 60 trials. Each trial started with the presentation of a monetary incentive (0.01€, 0.20€, 0.50€, 1.00€, 5.00€, 20.00€; shown in random order for 2-4 seconds), followed by a graduated force scale (4-6 seconds). The force scale ranged from 0 to 100%, with the top representing the maximum grip force of the individual. Visual feedback on the force scale indicated the current (grey bar) and peak in the current trial (red) grip force applied to the dynamometer. Reaching the top of this scale meant reaching one’s individual maximum force and thereby (A) winning the full incentive or (B) avoiding any loss. Each increment along the scale corresponded to a fraction of the monetary incentive. Afterwards, the win/loss and the current monetary total was displayed (3.5 seconds).

**Figure 1.**
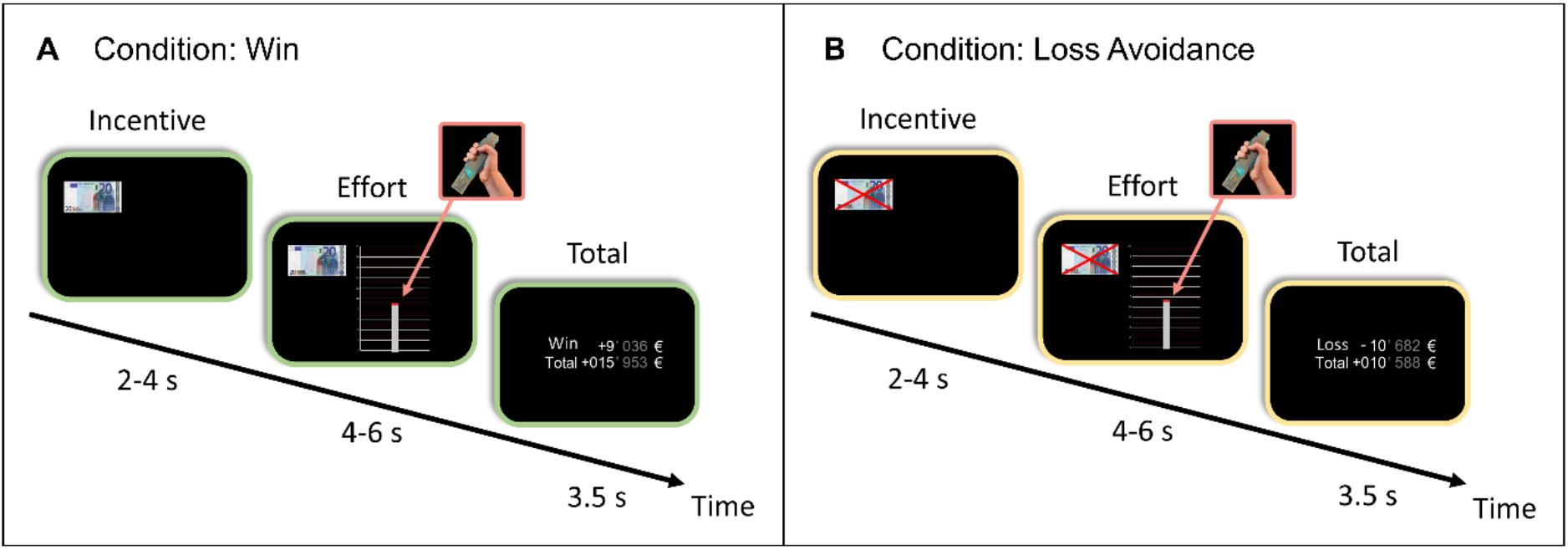
Behavioral incentivized force task. Illustration of one trial of each condition. (**A**) win and (**B**) loss avoidance. The red line moves conjointly with the gray bar upwards the scale when applying grip force to the dynamometer and represents the peak force within each trial.

### Data collection and quality checks

Grip force in the incentivized force task was sampled with 100 Hz. The peak force within the effort period (dependent variable) was extracted for each trial. MATLAB R19a was used to implement the task and to extract the dependent variable. Data were plotted and visually inspected for data quality purposes (i.e., ensuring that the whole force profile was captured) using Python 3.10.8.

### DaT-SPECT imaging

DaT SPECT images were obtained on a PRISM-3000 three-head SPECT system (Picker) following a standardized clinical procedure (i.e., [123I]Ioflupane injection, reconstruction using Chang’s attenuation correction, voxel-values normalized to the occipital cortex). Next, z-transformed deviation maps from age-matched healthy controls were computed using the EARL-BRASS software (Hermes, Sweden), and z-values of the bilateral putamen were extracted.^18^

### Statistical analyses

Using linear mixed models, we investigated the following effects on peak force: the interaction effect of task condition (win, loss avoidance) and group (HC, PD), the interaction effect of incentive level and group, and the interaction effect of putaminal dopaminergic terminal loss and incentive level in the PD group. Furthermore, to investigate the fatigue effect (decrease in force expenditure over time) we analyzed the interaction effect of trial number and group, and the interaction effect of putaminal dopaminergic terminal loss and trial number in the PD group. Incentive level was added to the model as categorical variable; and age, sex, and trial number (except for the fatigue analyses) were added as covariates. Additionally, all analyses were conducted with a second measure of force expenditure, replacing trial number with the cumulative area under the curve of the force-time curves. To account for repeated measures, participant ID was entered as a random factor. Type III Analysis of Variance Tables with Satterthwaite’s approximation of the degrees of freedom are reported and *p*-values < .05 were considered significant. Outliers were identified with the interquartile range method. All statistical analyses were conducted in R (version 4.1.2)^19^, linear mixed models were calculated with the R library lmerTest (version 3.1.3)^20^, and plots were created with the R library ggplot2 (version 3.4.0)^21^.

## Data availability

The data supporting this study’s findings are findable in the data registry of the CRC1451 (https://www.crc1451.uni-koeln.de/) and reasonable requests can be addressed to the corresponding author.

## Results

The HC and PD group did not significantly differ in age (HC: *M* = 63.75; PD: *M* = 62.4; *t*(38.13) = 0.61, *p* = .548), sex (*χ*^2^(1) = 0.74, *p* = .391) and MoCA (HC *Mdn* = 28, PD *Mdn* = 27), *W* = 312.5, *p* = .393). By design, patients with PD demonstrated a significantly higher UPDRS-III score (*Mdn* = 13) relative to HC (*Mdn* = 0), *W* = 3.5, *p* < .001.

843 out of 5640 trials were discarded from the analysis, as force profiles did not pass quality checks. We observed no significant difference between the two task-conditions (win vs. loss avoidance), *F*(1, 4747.2) = 0.52, *p* = .469. Thus, they were jointly analyzed. Analysis of the peak force revealed a significant interaction between group and incentive level, *F*(5, 4739.4) = 3.34, *p* = .005. Post hoc tests revealed that patients with PD applied significantly less force for the first four incentive levels (0.01€: *p* = .044, Δ = 7.63% of peak force; 0.20€: *p* = .006, Δ = 10.33% of peak force; 0.50€: *p* = .027, Δ = 8.37% of peak force; 1.00€: *p* = .018, Δ = 8.89% of peak force) compared to HC, but not for the highest two levels (5.00€: *p* = .06, Δ = 7.11% of peak force; 20.00€: *p* = .139, Δ = 5.6% of peak force; Fig. 2A). In the PD group, there was a significant interaction between putaminal dopaminergic terminal loss and incentive level, *F*(5, 2128.07) = 6.93, *p* < .001. Lower incentive levels were indeed associated with steeper reduction of peak force with putaminal dopaminergic terminal loss (Fig. 2B). The fatigue slopes of two subjects were identified as outliers and accordingly these subjects were excluded from the fatigue analyses. A significant interaction was observed between trial number and group, *F*(1, 4582) = 7.17, *p* = .007. Peak force of patients with PD decreased significantly faster across all 120 trials compared to HC (Fig. 2C). In the PD group, there was no significant interaction between trial number and putaminal dopaminergic terminal loss, *F*(1, 1970.30) = 0.73, *p* = .392 (Fig. 2D).

**Figure 2.**
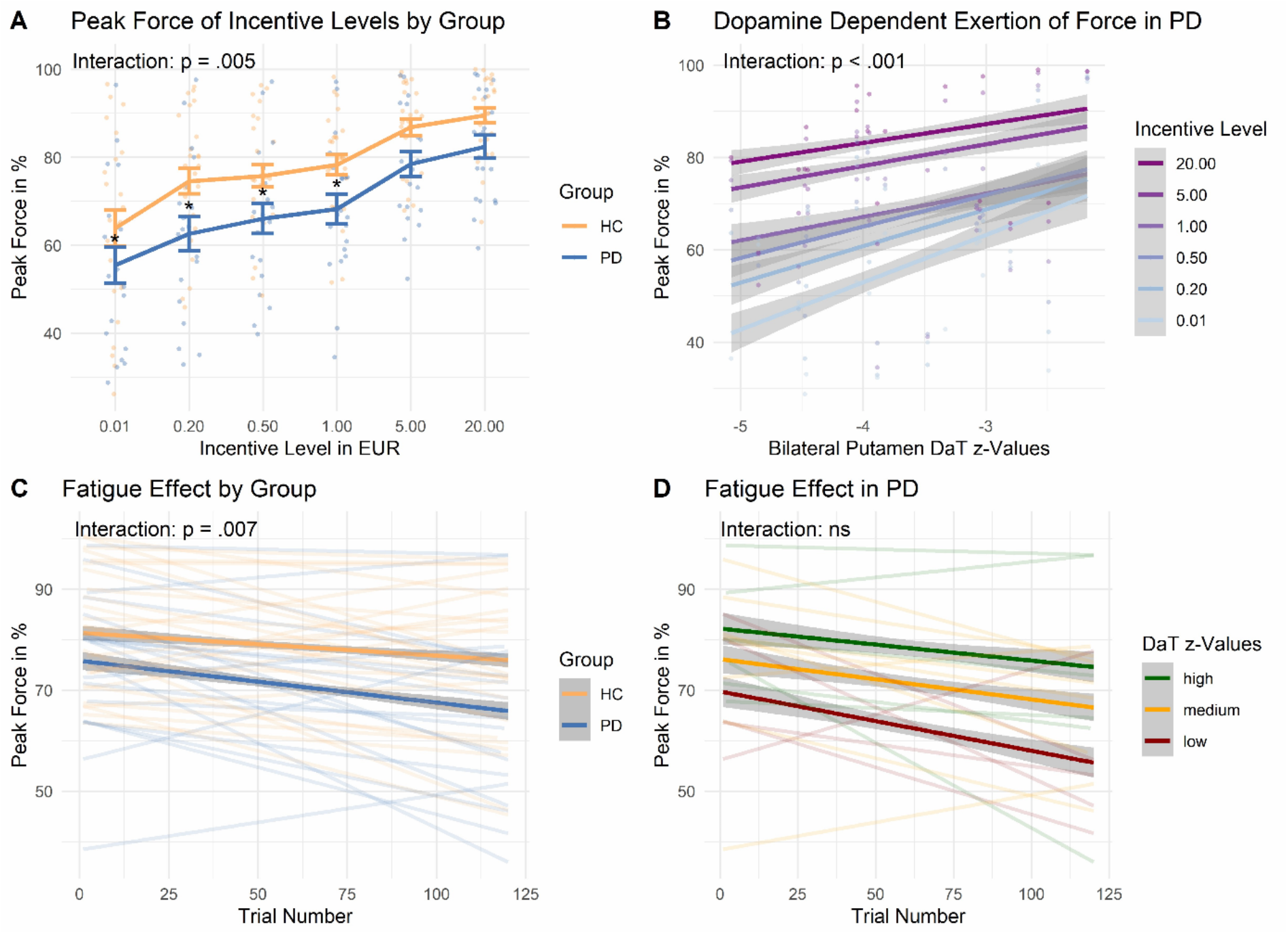
Results of the incentivized force task. **(A)** Peak grip force of patients with PD and HC for each incentive level. Standard error bars are depicted and significant group differences for incentive level 0.01, 0.20, 0.50, and 1.00 EUR are denoted by asterisks. (**B**) Correlation of peak grip force and putamen z-values of dopamine transporter density in the PD group for each incentive level. Lower incentive levels are associated with steeper slopes. (**C**) Peak grip force over all 120 trials of patients with PD and HC. The PD group demonstrates a significantly stronger fatigue effect (decrease in peak force). (**D**) Decrease in peak force over all trials is not associated with putaminal dopaminergic terminal loss in individuals with PD. ns = not significant

Analyses using the cumulative area under the curve revealed the same results, except for one post hoc tests of the significant interaction between group and incentive level. The lowest incentive level only demonstrated trend significance (0.01€: *p* = .051, Δ = 7.49% of peak force).

## Discussion

Here, we directly investigated if putaminal dopaminergic terminal loss in PD is associated with reduced motor effort in an incentivized force task. Our results demonstrate that patients with PD applied significantly less grip force than HC, especially for low incentive levels. Critically, this deficit was stronger for patients with a greater dopaminergic terminal loss.

Consistent with the motor motivation hypothesis, our findings indicate that patients with PD can attain physical performance levels comparable to those of healthy controls. However, this performance depended on the presence of high incentives, suggesting that patients normalized motor effort if the motivation is sufficiently high. This finding is in line with previous research, which demonstrated that patients with PD invested less effort compared to HC, but only for the lowest reward^13^. Moreover, it aligns with the phenomenon known as “paradoxical kinesis”, which describes an event where even severely immobilized patients with PD are able to move when much is at stake (e.g., running out of a burning house).^22–25^ Thus, the lack of movement in PD may be overcome through significant extrinsic motivators/incentives.

Importantly, dopamine loss seemed to decrease the motivation to exert physical effort. Previous studies have already indicated this by showing that patients with PD ON dopaminergic medication invest more effort compared to OFF medication.^13,26^ Thus, dopamine appears to increase the willingness to work for rewards. In line with previous findings, demonstrating the specificity of dopamine loss to low incentives,^13^ in our study the putamen dopamine transporter values were most strongly associated with effort in low incentive levels. Most likely, this means that the willingness to produce high effort is a function of both, incentive level (i.e., incentive-effort conversion rate) and putaminal dopamine level. In the absence of external incentives, the lack of dopamine takes greater effect. Alternatively, the statistical interaction effect could have been produced by a ceiling effect with regard to motor effort. In other words, the cumulative effect of dopamine and external incentives could be “maxed out” at around 90% peak force. An interesting prospect for future studies could be the question whether a greater deficit will also affect high incentives or leave them intact because they are less dopamine dependent. In this context, we want to emphasize that our patient cohort consisted of individuals with only mild motor symptoms.

We found that patients with PD not only demonstrated grip force which was lower on average, but which also decreased faster over all trials, denoting a greater exhaustion over time, which has also been demonstrated by previous research.^27^ Furthermore, based on existing studies and the present findings we expected greater putaminal dopaminergic terminal loss to be linked to a greater fatigue effect. Notably, this was not the case as decrease in force expenditure was not associated with the putaminal dopaminergic terminal loss (Fig. 2D). Even though a greater dopaminergic deficit was associated with lower force in general, it did not affect the force expenditure over time. Thus, the effect of dopamine depletion to reduce movement vigor seems to be stable over time periods of physical exertion.

A few potential shortcomings need to be considered. First, given that our findings are based on a limited number of subjects, the results should be interpreted with caution. Furthermore, we only had DaT SPECT imaging data of the PD group and thus, some of the analyses were not feasible in HC.

Taken together, our study provides important evidence for a critical role of dopamine in motivating movements. We found that patients with PD demonstrate decreased motivation for effortful behavior if incentives are low, suggesting that dopamine is critical to overcome effort when the trade-off with the benefit is poor. Moreover, for the first time, we can show that this decrease in effort is quantitatively associated with the degree of dopamine depletion in the putamen, as especially manifest in low reward conditions, pointing to a progressive change with dopamine loss severity in the cost-benefit computation of lower yield movements.

